# Functional wiring of the human medial temporal lobe

**DOI:** 10.1101/257899

**Authors:** E. A. Solomon, J. M. Stein, S. Das, R. Gorniak, M. R. Sperling, G. Worrell, C. Inman, B. Lega, B. C. Jobst, D. S. Rizzuto, M. J. Kahana

**Author notes:** Corresponding author: Michael J. Kahana, 425 S. University Ave. Levin Building, Rm. 201, Philadelphia, PA 19104.

## Abstract

The medial temporal lobe (MTL) is a locus of episodic memory in the human brain. It is comprised of cytologically distinct subregions that, in concert, give rise to successful encoding and retrieval of context-dependent memories. However, the functional connections between these subregions are poorly understood. To determine functional connectivity among MTL subregions, we had 126 subjects fitted with indwelling electrodes perform a verbal memory task, and asked how encoding or retrieval correlated with interregional synchronization. Using phase-based measures of connectivity, we found that synchronous theta (4-8 Hz) activity underlies successful episodic memory, whereas high-frequencies exhibit desynchronization. Moreover, theta functional connectivity during encoding aligned with key anatomic connections, including critical links between the entorhinal cortex, dentate gyrus, and CA1 of the hippocampus. Retrieval-associated networks demonstrated enhanced involvement of the subiculum, reflecting a substantial reorganization of the encoding-associated network. We posit that coherent theta activity within the MTL marks periods of successful memory, but distinct patterns of connectivity dissociate key stages of memory processing.

**Significance Statement:** The brain functions through the interaction of its distinct parts, but little is known about how such connectivity dynamics relate to learning and memory. We used a large dataset of 126 human subjects with intracranial electrodes to assess patterns of electrical connectivity within the medial temporal lobe – a key region for memory processing – as they performed a memory task. We discovered that unique networks of time-varying, low-frequency interactions correlate with memory encoding and retrieval, specifically in the theta band. Simultaneously, we observed elevated spectral power at high frequencies in these same regions. The result is a complete map of physiological dynamics within the MTL, highlighting how a reorganization of theta networks support distinct memory operations.

## Introduction

Storing episodic memory is an inherently integrative process, long conceptualized as a process that links information about new items to an observer’s current thoughts, emotions, and environment (1). Decades of behavioral observations, clinical case studies, and scalp electroencephalography (EEG) in humans have shed light on the key principles and diverse set of brain structures underlying this integration, including frontal, lateral temporal, and medial temporal cortex (MTL) (2). Recent hypotheses invoke the idea that communication among these regions supports memory formation, spurred by a growing number of functional imaging and intracranial EEG (iEEG) experiments that show synchronized activity among the MTL and cortical structures during memory tasks (3–6).

However, the MTL has a unique role in supporting episodic memory. Damage to the MTL results in profound deficits of memory (7), and it has been shown to exhibit enhanced neural activity during memory processing in a range of tasks and experimental models (8, 9), identifying this area as a key anatomic hub of episodic encoding and retrieval. The MTL is structurally complex; it is subdivided into hippocampus, rhinal cortex, and parahippocampal cortex. The cornu ammonis (CA), dentate gyrus, and subiculum comprise the hippocampus, while the entrorhinal and perirhinal cortices form rhinal cortex. Microscale recordings in animals have revealed that these substructures exhibit distinct patterns of activity during memory and navigation tasks, including the generation of oscillations, inter-regional synchronization, and neuronal selectivity for time and space (10–16). Computational models of MTL function have assigned unique roles to MTL substructures, pertaining to episodic encoding, retrieval, or recognition (10, 17–20) – typically, these models suggest extrahippocampal regions are responsible for placing sensory inputs in a useful representational space, while the hippocampus itself forms associative links between these representations and their prevailing context.

Notably, virtually all of the aforementioned animal and modeling literature suggests that MTL substructures communicate with one another as they engage in memory processing. However, the volume of aforementioned work on intra-MTL connectivity has not been matched by validation studies in humans. Though a handful of investigations have begun to address this question in neurosurgical patients (21–23), limited electrode coverage and imprecise localizations have made it difficult to study the fine spatial scale and complete extent of neural synchronization within the MTL. But doing so is a crucial component of (1) confirming that intra-MTL synchronization observed in animal models also correlates with memory processing in humans, and (2) validating models of MTL function that suggest communication among specific regions – such as rhinal cortex versus hippocampus – supports computations necessary for associative memory formation and retrieval.

The use of intracranial depth electrodes to study neural activity in the MTL also allows neural activity to be studied at different timescales. Slow theta (4-8 Hz) oscillations in the hippocampus have been observed during memory processing in humans (24–27), as have fluctuations at higher frequencies, including the gamma (30-60 Hz) band (28, 29). These oscillations have been theorized to support synchronization between neural assemblies in the MTL (15, 17, 30–32), but MTL connectivity has not been fully mapped across frequency bands. The extent to which different frequencies underlie neural synchronization in memory therefore remains an open question, though converging lines of evidence strongly suggest the most prominent connectivity effects occur at low frequencies (33–37). Moreover, no studies have deeply considered how observations of within-MTL synchronization reflect an experimenter’s choice of connectivity metric, including those that are designed to limit the effects of volume-conducted signals that may affect connectivities measured across relatively short distances (38).

In this study, we aimed to define the patterns of functional connectivity that emerge in the human MTL and to specifically characterize how MTL-subregional connectivity differs when memories are being stored versus when they are being subsequently retrieved. Additionally, we asked whether functional networks were sensitive to the choice of connectivity metric, utilizing the phase-locking value (PLV) and weighted phase-lag index (wPLI; insensitive to volume conduction). We leveraged a large dataset of 126 subjects with depth electrodes placed in the MTL, localized with hippocampal subfield resolution, and focused on two key contrasts: (1) the encoding subsequent memory effect (SME), differentiating remembered from forgotten items, and (2) successful retrieval versus periods of unsuccessful memory search. We found that successful encoding was characterized by low-frequency connections which converged on left entorhinal cortex, while retrieval was associated with enhanced theta connectivity to the subiculum and CA1. However, these differing connectivity patterns were not correlated with markedly different patterns of local spectral power between encoding and retrieval, suggesting functional connections are a key mechanism by which the MTL may switch between distinct memory operations. Furthermore, we noted that theta-band connectivity was present regardless of choice of connectivity metric or referencing scheme, though connectivity was generally blunted when using the wPLI. Taken together, our findings show that low-frequency functional coupling in the MTL supports the formation of new memories, with the specific pattern of connections acting as the key determinant of successful encoding and retrieval, respectively.

## Results

Our general approach to characterizing intra-MTL connectivity was to (1) examine the structure of functional connectivity networks using graph-theoretic analysis, (2) examine the timecourse of connectivity in key connections, and (3) relate changes in connectivity to changes in local activity, as reflected by spectral power. To do this, we correlated intra-MTL synchronization with memory state using two contrasts in a verbal free-recall paradigm. First, we examined the subsequent memory effect (SME), which has been widely employed to characterize whole-brain modulations of spectral power (e.g. Burke et al., 2014a, 2014b) that correlate with successful memory encoding. Second, we examined a memory retrieval contrast, wherein epochs of time leading up to verbalization of a recalled item are compared to matched epochs of time, from other word lists, where no recall occurs (e.g. 29, 40, 41; see Methods for details). We refer to these matched periods as “deliberation” intervals.

For each contrast, we constructed intra-MTL functional connectivity maps at each frequency band, using the phase-locking value (PLV) and weighted phase-lag index (wPLI). The PLV (42) has long been used to assess whether, across trials, there is a consistent phase difference between two electrodes. The wPLI is a more recent modification of the PLV (38), which downweights phase differences near zero under the assumption that two such electrodes are detecting a volume-conducted signal through brain tissue, and not true physiologic coupling (Figure 1D). We had 126 subjects perform a verbal free-recall task during which iEEG was collected from depth electrodes placed in the MTL. Subjects were serially presented with 12-item word lists and asked to recall as many words as possible after a brief distractor task (Figure 1A-C; see Methods for details). For each electrode pair, phase differences were computed for each trial, i.e. an encoding or retrieval event. Trials were sorted by whether a word was later recalled or forgotten (or, in the retrieval contrast, a successful retrieval event or matched deliberation; Figure 1E). PLV/wPLI were computed for successful/unsuccessful groups separately, and tested for significant differences via a nonparametric permutation procedure (see Methods for details). Effects were averaged across electrode pairs, subjects, and time, yielding a z-score that indicates the relative synchronization in successful vs. unsuccessful memory encoding/retrieval for each pair of MTL regions (see Figure 2A for an example pair; see Supplemental Figure 1 for subject count per region-pair). Unless indicated otherwise, we use a common average reference for all analyses; the bipolar reference is used in some cases to demonstrate generality of key results regardless of referencing scheme.

**Figure 1.**
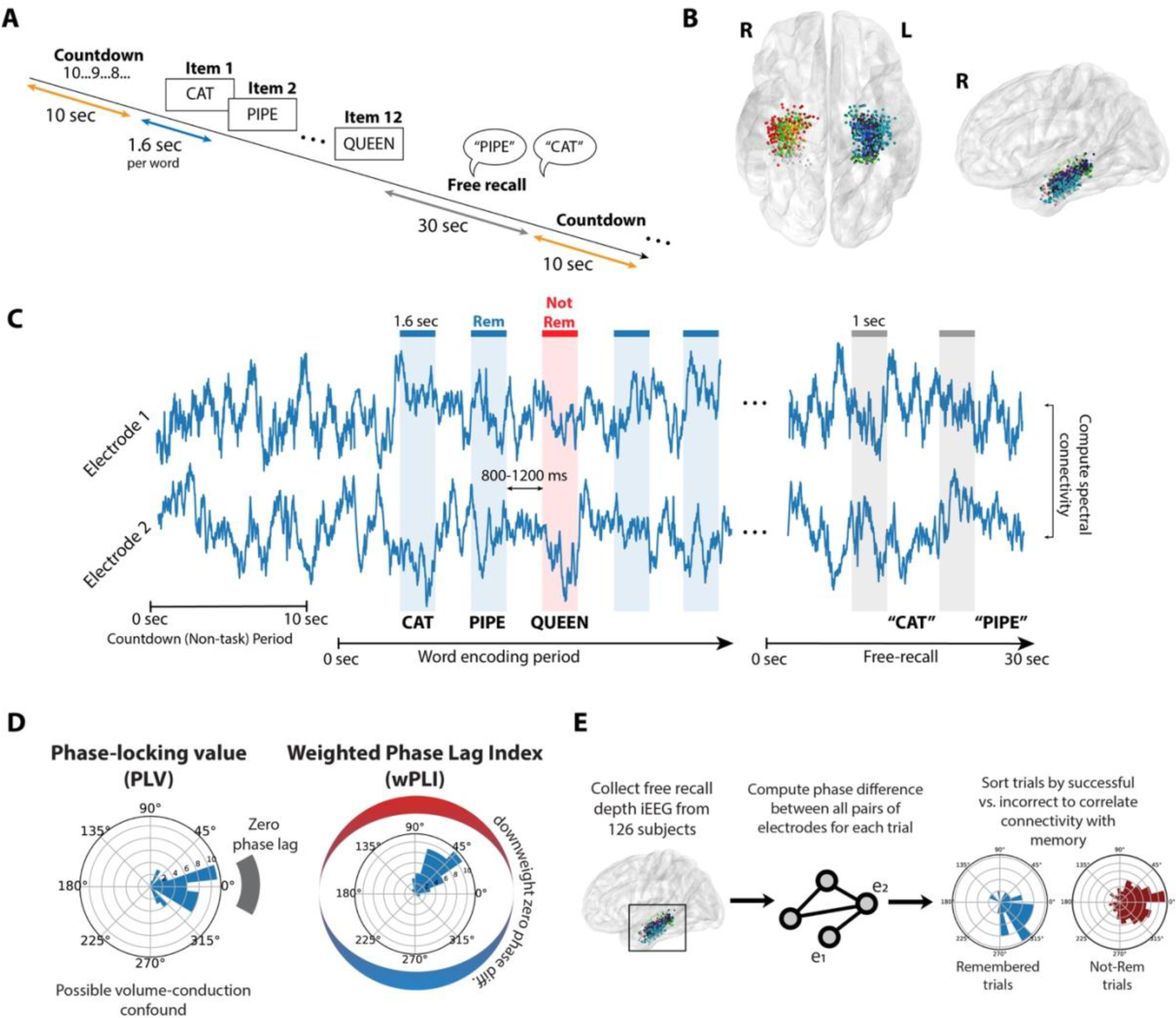
Task structure and analysis methods. **A.** Subjects performed a verbal free-recall task, consisting of alternating periods of pre-list countdowns (orange), word encoding (blue), and free-recall (gray). See Methods for details. **B.** 126 subjects with indwelling electrodes in the medial temporal lobe (MTL) participated. Electrodes were localized to CA1, CA3, dentate gyrus (DG), subiculum (Sub), perirhinal cortex (PRC), entorhinal cortex (EC), or parahippocampal cortex (PHC). Each dot shows an electrode in this dataset, colored by MTL subregion. **C.** To construct networks of intra-MTL activity, we used PLV and wPLI to analyze phase differences between electrode pairs. Time windows in two conditions were analyzed: 1.6-second epochs during word encoding (blue/red), and 1-second periods leading up to recall vocalizations (gray). **D.** PLV reflects the consistency of phase differences across trials, indicated in example data as clustered phase lags around zero (left). The wPLI works similarly, but downweights the contribution of phase lags near the zero axis, which may reflect volume conduction (right). Accordingly, stronger connectivity is observed if phase lags are clustered around a direction far from zero (or 180 degrees). **E.** To assess intra-MTL connectivity, phase differences are computed for each electrode pair in all trials, and trials are then sorted by successful vs. unsuccessful memory. PLV or wPLI is computed for each distribution, and a nonparametric permutation procedure is used to determine whether connectivity is significantly different between distributions. Connectivity values are averaged across electrode pairs and subjects to yield the final MTL network maps depicted in Figure 2 (see Methods for details).

**Figure 2.**
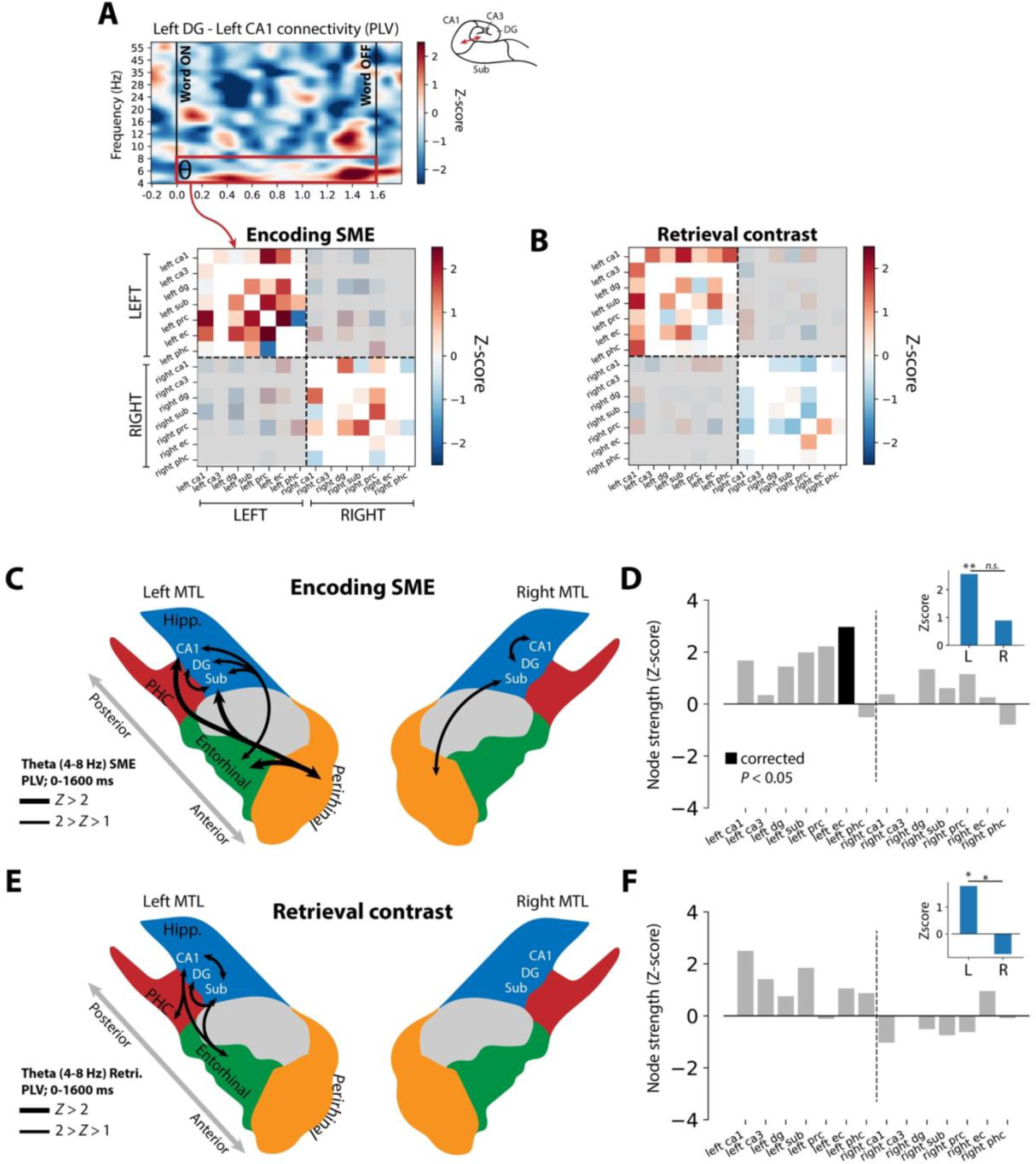
Structure of theta networks supporting episodic memory. **A.** To determine overall connectivity for each pair of MTL subregions, PLV or wPLI is averaged over the encoding (word presentation, 0-1.6 seconds) or retrieval (−1.0 to 0 seconds prior to retrieval onset) intervals, yielding a single z-scored connection weight. (see Methods for details). The matrix representation of all these weights is called an adjacency matrix, shown here for the encoding contrast in the theta band (PLV). Any inter-regional connection with fewer than 5 subjects’ worth of data is excluded from analysis (white cells). Because interhemispheric connections are less well sampled than intra-hemispheric connections, and because interhemispheric connectivity is largely asynchronous, they are excluded from this analysis of network structure (gray shading). **B.** Retrieval contrast theta adjacency matrix (PLV), organized as in (A). **C.** Depiction of strongest (Z > 1) synchronous PLV connectivity in the SME contrast, derived from the theta adjacency matrix in (A). These connections reflect the averaged connection strength over the word presentation interval (0.0-1.6 seconds; see Methods for details). Thicker lines reflect Z-scores above 2. **D.** Z-scored node strength for each MTL region, computed only for connections to ipsilateral MTL regions (see Methods for details). Node strength indicates the sum of all connections to a given region, with positive Z-scores indicating enhanced overall connectivity to a given region during successful encoding epochs (a “hub” of connectivity). Left EC exhibited significant positive node strength (FDR-corrected permuted *P* < 0.05) correlated with words that were successfully remembered. **Inset:** Z-scored total network strength for all intra-hemispheric MTL connections, computed by summing the connection weights for each hemisphere’s MTL subregions separately. Intra-MTL connections on the left are significantly greater than chance (*P* = 0.005), and trend greater than right-sided connections (*P* = 0.15). **E.** Schematic of strongest theta retrieval connections, reflecting increased PLV between two MTL subregions in the 1-second immediately prior to successful retrieval of a word item. **F.** Same as (D), but reflecting synchronous activity from the 1-second period prior to successful retrieval of a word item. No region exhibits a significant node strength after correction for multiple comparisons, but left CA1 is significant if uncorrected (*P* = 0.013). **Inset:** Z-scored total network strength for all intra-hemispheric MTL connections. Left-sided connections are significantly greater than chance (*P* = 0.04) and significantly greater than right-sided connections (*P* = 0.03).

### Theta networks of memory encoding and retrieval

Given strong evidence in the literature for synchronous memory effects in the theta band (33–37), we first sought to characterize the detailed structure of theta networks in the MTL. To do this, we asked whether any regions acted as “hubs” of the MTL by computing the node strength for each region, using theta PLV connection weights. Node strength reflects the overall connectivity to a given node of the network by summing all of its connection weights. In the SME contrast, left entorhinal cortex emerged as a significant hub (corrected permuted *P* < 0.05; Figure 2D). In the retrieval contrast, left CA1 was numerically greatest and significant if not corrected for multiple comparisons (permuted *P* = 0.013; Figure 2F). The single strongest connection for the encoding/retrieval contrasts were EC-PRC (*Z* = 2.65) and CA1-Sub (*Z* = 2.00), respectively (see Figure 1 legend for region abbreviations). For each contrast, the strongest synchronous connections are depicted schematically in Figures 2C and 2E. In both retrieval and encoding, entorhinal cortex exhibits enhanced connectivity to CA1 and subiculum, with additional perirhinal-hippocampal connections present exclusively in encoding. Additionally, in both contrasts, connections within the left MTL are significantly greater than zero (encoding, *P* = 0.005; retrieval, *P* = 0.04), and stronger than connections within the right MTL, though not significantly so for encoding (encoding, *P* = 0.15; retrieval, *P* = 0.03).

These findings align with known anatomical connections and functional roles of MTL subregions. The entorhinal cortex acts as a key input structure to the hippocampus and represents the convergence of information from the perirhinal and parahippocampal cortices (45, 46) – it is fitting that this structure exhibits enhanced theta connectivity to other MTL structures during with successful memory encoding. Furthermore, a reorganization of theta networks featuring enhanced connectivity between the subiculum and CA1 comports with anatomical connectivity and notion of subiculum’s role as a major output structure of the hippocampus (47, 48). However, this network-based analysis (1) averages synchrony effects over the entire word presentation or retrieval intervals, obscuring time-varying dynamics, and (2) considered only PLV connectivity, which may reflect nonphysiologic volume conduction.

### Temporal dynamics of memory-related connectivity

Having shown that encoding- and retrieval-associated theta networks differ in their structure but align with known anatomical connectivity of the MTL, we next asked whether our previously-identified synchronous connections exhibited time-varying dynamics. Additionally, we aimed to determine whether these connectivity effects were robust to use of the PLV or wPLI, which discounts phase differences near zero. In particular, we hypothesized that two possibilities could underlie changes in PLV and wPLI between conditions. First, tightly-clustered phase lags in a nonzero direction could rotate towards zero, resulting in a reduced wPLI since phase lags are downweighted closer to zero (Figure 3A, top row). In this case, PLV would remain unchanged. However, the correction for volume-conduction comes at a cost: tightly-clustered phase lags could still be well within a believable, nonzero range, yet wPLI would indicate a relative decrease in synchronization between conditions. The second possibility is that tightly clustered nonzero phase lags could increase in variance between conditions, resulting in decreased PLV and wPLI (Figure 3A, bottom row). In experimental reality, both between-condition changes could occur simultaneously.

**Figure 3.**
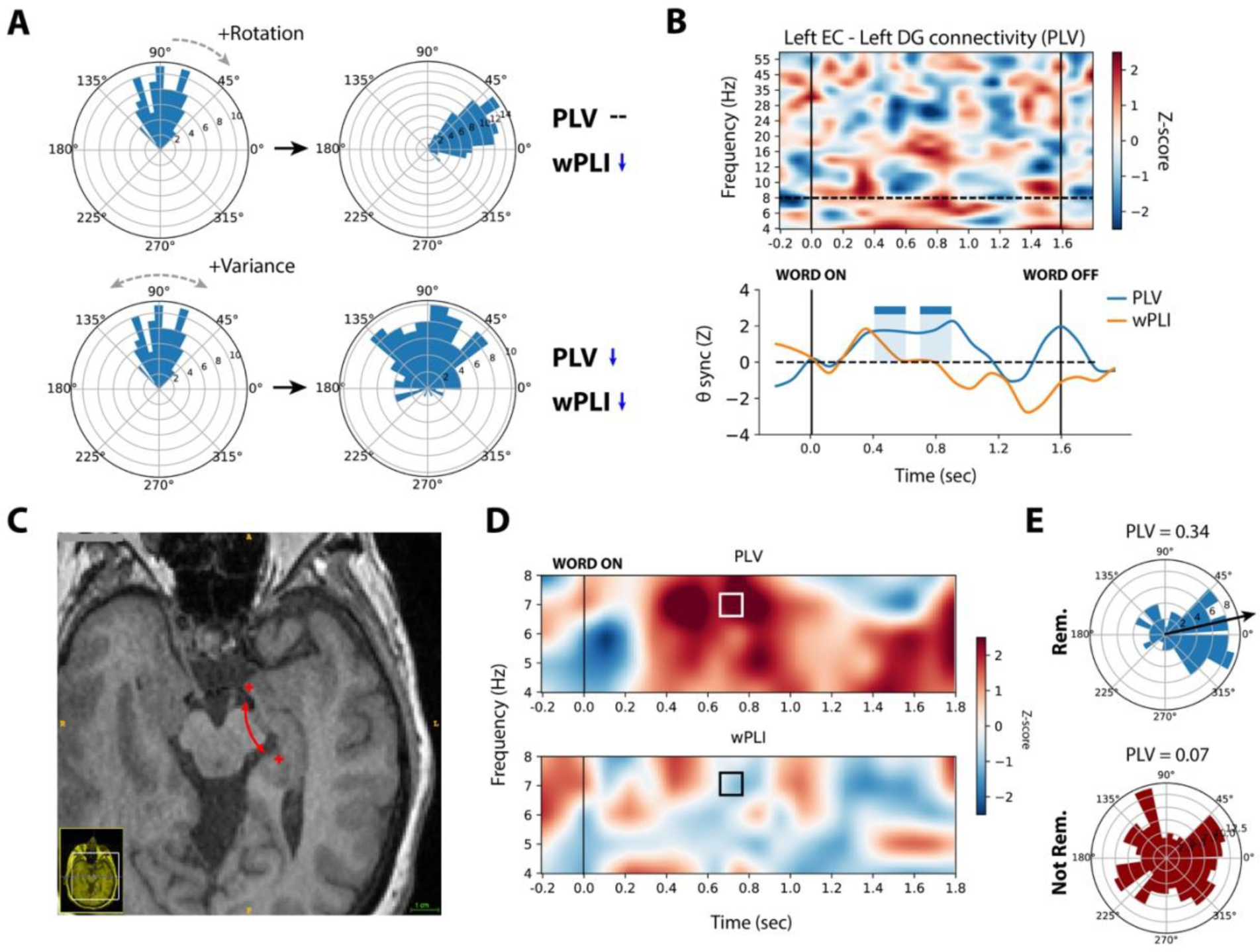
Time-varying dynamics of left EC to left DG coupling. **A.** As demonstrated using simulated data, the wPLI can differ between conditions for two reasons. First, depicted in the top row, well-clustered phase lags can rotate toward zero – decreasing wPLI which downweights phase lags closer to zero to account for volume conduction. In this case, the PLV will remain unchanged. Second, depicted in the bottom row, the variance of a nonzero phase lag distribution can increase between conditions, causing decreases in PLV and wPLI. **B.** *Top:* Time-frequency plot of PLV synchronization between left EC and left DG in the encoding contrast, averaged across subjects. Red colors indicate relative synchronization in remembered versus not-remembered trials, blue colors indicate relative desynchronization. Vertical black lines indicate word onset and offset, dotted horizontal line indicates the max theta frequency considered for connectivity analyses. *Bottom:* In non-overlapping 100 ms windows, PLV (blue) and wPLI (orange) values are averaged across all theta frequencies, yielding a synchronization timeseries. Any two consecutive 100 ms windows with synchronization significantly greater than chance (*P* < 0.05) are marked as significant with blue or orange colored rectangles (see Methods for details). In this region-pair, PLV is significant from 400-600 ms and 700-900 ms after word onset. **C.** Axial T1-weighted MRI slice depicting electrode locations for a representative pair spanning the left DG and EC. **D.** Time-frequency plots for the indicated electrode pair, theta frequencies only. Top row shows PLV synchronization during successful encoding, bottom row shows wPLI synchronization. The considered interval and frequency for phase lags in (E) is marked with a square. **E.** For the highlighted frequency and interval, phase lags were binned according to whether the trial was later remembered (blue) or not-remembered (red). The mean direction of the clustered remembered trials is 9.7 degrees (PLV = 0.34), indicated with the black arrow. Not-remembered trials are unclustered, as reflected by a low PLV of 0.07.

To illustrate this, we examined phase lag distributions for a key connection in the encoding contrast – left EC versus left DG. Across all subjects, we observed significantly enhanced theta PLV (permuted *P* < 0.05 from 400-600 ms and 700-900 ms) with successful encoding, though no significant wPLI during those same intervals (Figure 3B). In a representative subject who exhibited enhanced PLV and minimal wPLI modulation in similar intervals, we examined phase lags between remembered and not-remembered trials for a pair of electrodes in the left EC and DG (Figure 3C-D). Phase lags for remembered trials were clustered near zero (PLV = 0.34, mean direction = 9.7 degrees), and uniformly distributed in the not-remembered condition (PLV = 0.07; Figure 3E). One interpretation of these results is that wPLI is operating as intended; small phase lags could be reflecting volume-conducted signal from a common source that should be discounted. Therefore, even the tightly-clustered phase distribution in the remembered condition does not yield a relative increase in wPLI. Conversely, it is not clear whether a mean phase lag of 9.7 degrees is close enough to zero to justify substantially discounting those signals; prior studies have excluded phase differences less than 5 degrees from zero (33). Finally, in this example, the DG and EC recording contacts are separated by more than 2 cm, a spacing which is greater than putative distances in which common signal is conducted in brain tissue (49). It is therefore possible that use of wPLI is inappropriately rejecting true, near-zero phase coupling.

Temporal dynamics of theta connectivity differed between MTL subregion pairs. Left EC and left PRC showed significantly enhanced PLV connectivity (permuted *P* < 0.05) in the −100-300 ms interval relative to word onset (Figure 4A). Left EC and left CA1, linked by the perforant pathway, exhibited enhanced PLV synchrony from 500 ms to 800 ms after word onset, similar to the EC-DG connection (Figure 4B). In both cases we observed subthreshold increases in wPLI during the word encoding interval.

**Figure 4.**
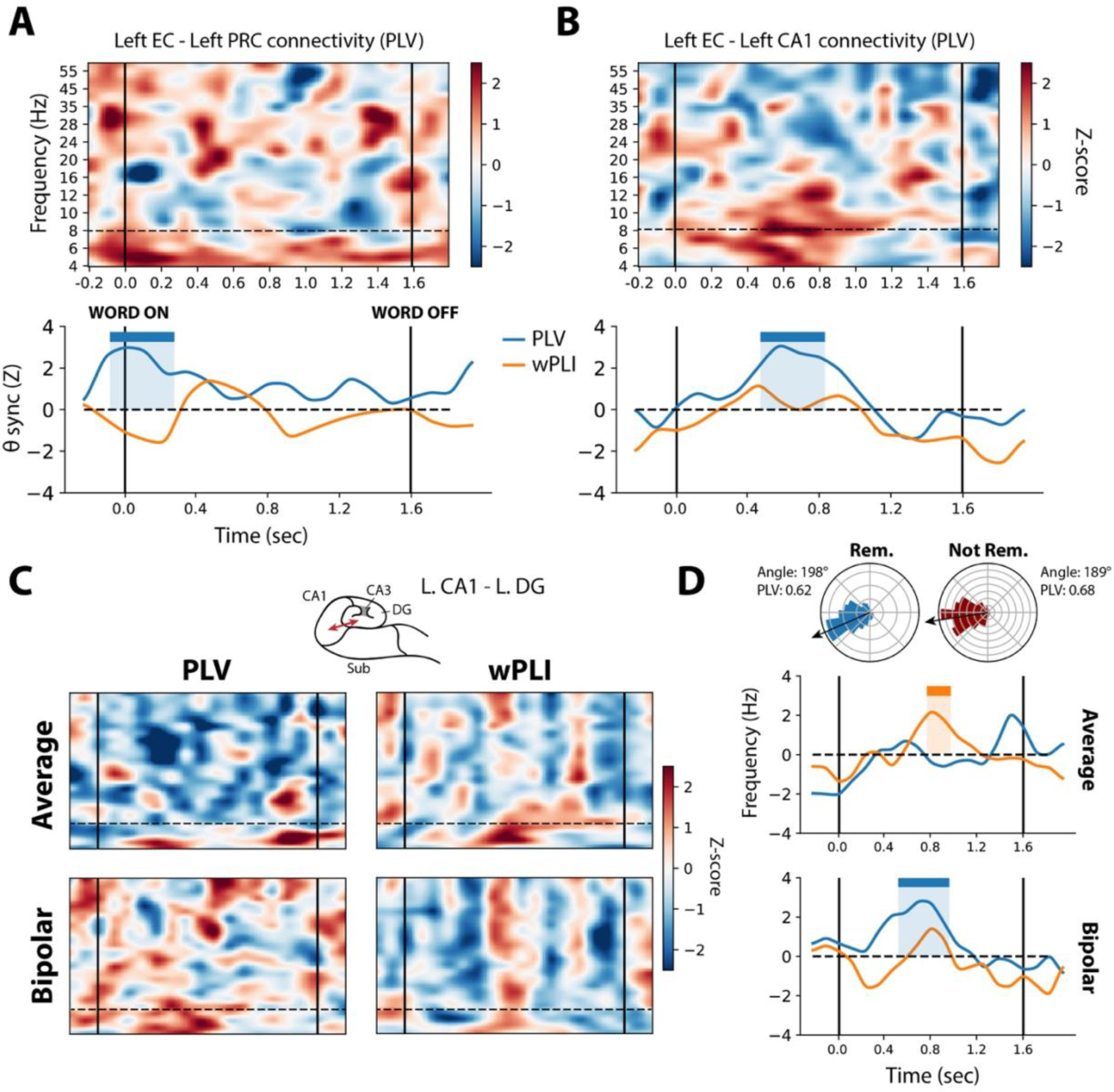
Timing analysis of key encoding connections. **A.** *Top:* Time-frequency plot of left EC-PRC synchronization (PLV) in the encoding contrast, averaged across subjects. *Bottom:* Timecourse of PLV/wPLI theta synchronization, averaged across subjects. Significant (*P* < 0.05) PLV synchronization is marked from −100 to 300 ms after word onset. **B.** Same as (A), but for the EC vs. CA1 connection. Significant PLV synchronization is marked from 500 to 800 ms after word onset. **C.** *Top row:* PLV and wPLI time-frequency plots for left CA1 vs. left DG, under the common average reference. *Bottom row:* PLV/wPLI time-frequency plots under the bipolar reference. See Methods for details. **D.** For the left CA1 vs. left DG connection, timecourses of theta PLV/wPLI connections, organized as in (A). Significant wPLI was observed under the average reference from 800-1000 ms, while significant PLV was observed under the bipolar reference from 500-900 ms. Phase lag distributions from a representative electrode pair (average reference) are depicted above the timeseries, indicating a relative rotation away from zero for remembered trials.

Connectivity between left CA1 and left DG illustrates convergent results regardless of connectivity metric or referencing scheme. We observed general increases in theta connectivity in all measured conditions (Figure 4C), though PLV increases were subthreshold under the average reference, while wPLI increases were significant (permuted *P* < 0.05, 800-1000 ms; Figure 4D). To understand this unique case, we examined the phase lag distributions for a representative electrode pair (Figure 4D, top row). In both remembered and not-remembered distributions, phase lags were tightly clustered (PLV = 0.62, 0.68 respectively), though the not-remembered distribution was rotated towards the zero axis (−9 degrees versus −18 degrees). This rotation yielded a relative increase in wPLI even as overall phase lag clustering fell slightly. Notably, we also found enhanced PLV in this region-pair under the bipolar reference (500-900 ms) and a concomitant subthreshold wPLI increase (Figure 4D, bottom panel).

Finally, synchronization during memory retrieval intervals also exhibited time-varying structure. In two key connections, left CA1 vs. subiculum and left CA1 vs. left EC, we observed increases in theta connectivity in the period leading up to vocalization of a recalled word (CA1-Sub, *P* < 0.05 −200-0 ms prior to retrieval; CA1-EC, −300-100 ms; Figure 5A-B). In a representative electrode pair, we noted CA1-Sub synchronization was associated with clustered near-zero phase lags (PLV = 0.47, 8.6 degrees; Figure 5C left), explaining no observed increase in relative wPLI. In the CA1-EC pair, phase lags were well-clustered in both conditions but rotated near zero for successfully retrieved events (PLV = 0.70, 1.9 degrees; Figure 5C right).

**Figure 5.**
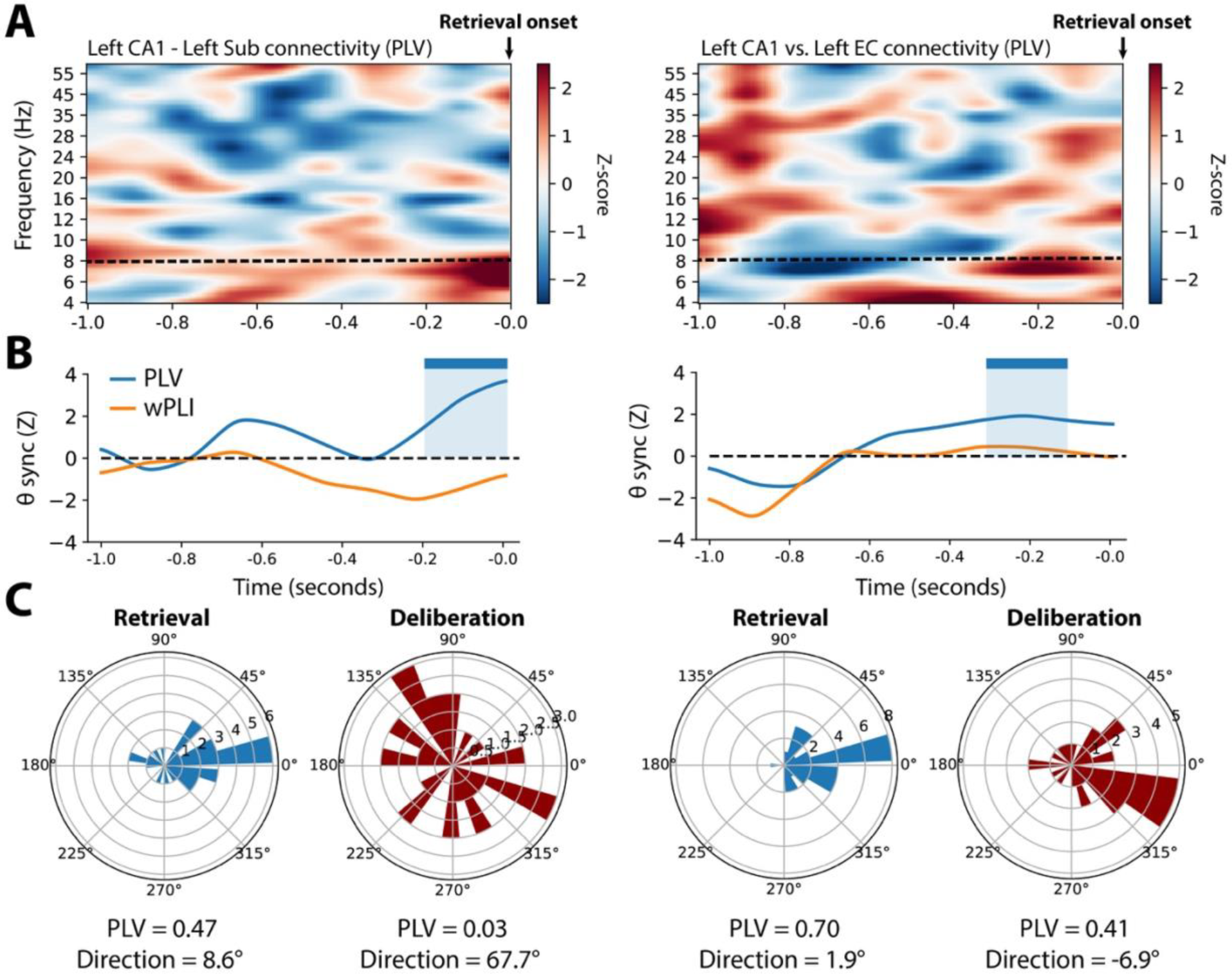
Timing analysis of key retrieval connections. **A.** *Left:* Time-frequency plot of left CAl-subiculum PLV synchronization in the retrieval contrast, averaged across subjects. *Right:* Time-frequency plot of left CA1-EC PLV synchronization in the retrieval contrast. **B.** *Left:* Timecourse of left CA1-subiculum PLV/wPLI synchronization, organized as in Figure 4B. Significant (*P* < 0.05) PLV synchronization is marked from −200 to 0 ms prior to recall onset. *Right:* Timecourse of left CA1-EC synchronization. Significant PLV synchronization is marked from −300 to −100 prior to recall onset. **C.** *Left:* In a representative CA1-sub electrode pair demonstrating enhanced PLV in the significant interval, successfully-retrieved trials demonstrated phase lag clustering around a mean direction of 8.6 degrees (PLV = 0.47). No clustering was evident for epochs where no recall occurred, called “deliberation” trials. *Right:* In a representative CA1-EC electrode pair, successful retrieval events showed greater phase clustering than deliberation events, but retrieval events were aligned with the zero axis (mean direction, 1.9 degrees). Therefore, PLV reflected a memory-related increase, but wPLI did not.

Taken together, we found that the general increase in left-MTL synchronization is driven by time-varying increases in connectivity between key regions, including left EC, CA1, DG, and subiculum. Connections to or within the hippocampus typically occurred more than 500 ms after word onset in the encoding interval, while EC and PRC exhibited an early synchronization approximately coincident with word onset. Differences between observed PLV and wPLI derive from complex differences in underlying phase lags, though we note that clustered phase lags near – but not at – zero tend to blunt wPLI’s sensitivity to memory-related effects.

### Relationship between connectivity and spectral power

Our primary focus was to characterize patterns of intra-MTL connectivity, but it is known that MTL subregions exhibit distinct patterns of local activation associated with episodic memory (e.g. 50, 51). We therefore asked whether changes in local spectral activity within the MTL correlate with encoding and retrieval states, and whether such changes relate to inter-regional theta connectivity. To do this, we analyzed the relative spectral power between successful and unsuccessful encoding/retrieval trials, in the theta band (4-8 Hz) and frequencies that correspond to high-frequency activity (HFA, 30-90 Hz). HFA is established as a general marker of neural activation that likely includes gamma oscillatory components and spectral leakage from aggregate unit spiking activity (52). For each MTL subregion, we computed the power SME and retrieval contrast for each electrode at each frequency, and averaged these effects across electrodes and subjects (see Methods for details). This procedure results in a t-statistic that reflects the relative power in a given region between successful and unsuccessful encoding/retrieval events.

Though we broadly observed positive theta connectivity associated with successful episodic memory, spectral power contrasts at the same frequencies went in the opposite direction. Bilateral CA1 and PRC exhibited significant decreases in theta power associated with successful encoding, as did left DG, left PHC, and right subiculum (FDR-corrected *P* < 0.05; Figure 6A). Bilateral CA1 also exhibited significantly enhanced HFA, and HFA was otherwise nonsignificantly increased in all MTL regions. Power dynamics associated with successful retrieval were similar to those observed in the encoding contrast. Theta was generally decreased in the left MTL, significantly so in left PRC and CA1 (FDR-corrected *P* < 0.05). Furthermore, HFA was elevated in bilateral CA1 and DG. The general trend of decreased theta power and increased HFA aligns with a robust literature demonstrating this same effect across a diverse array of cortical regions and the MTL (3, 28, 29, 53).

**Figure 6.**
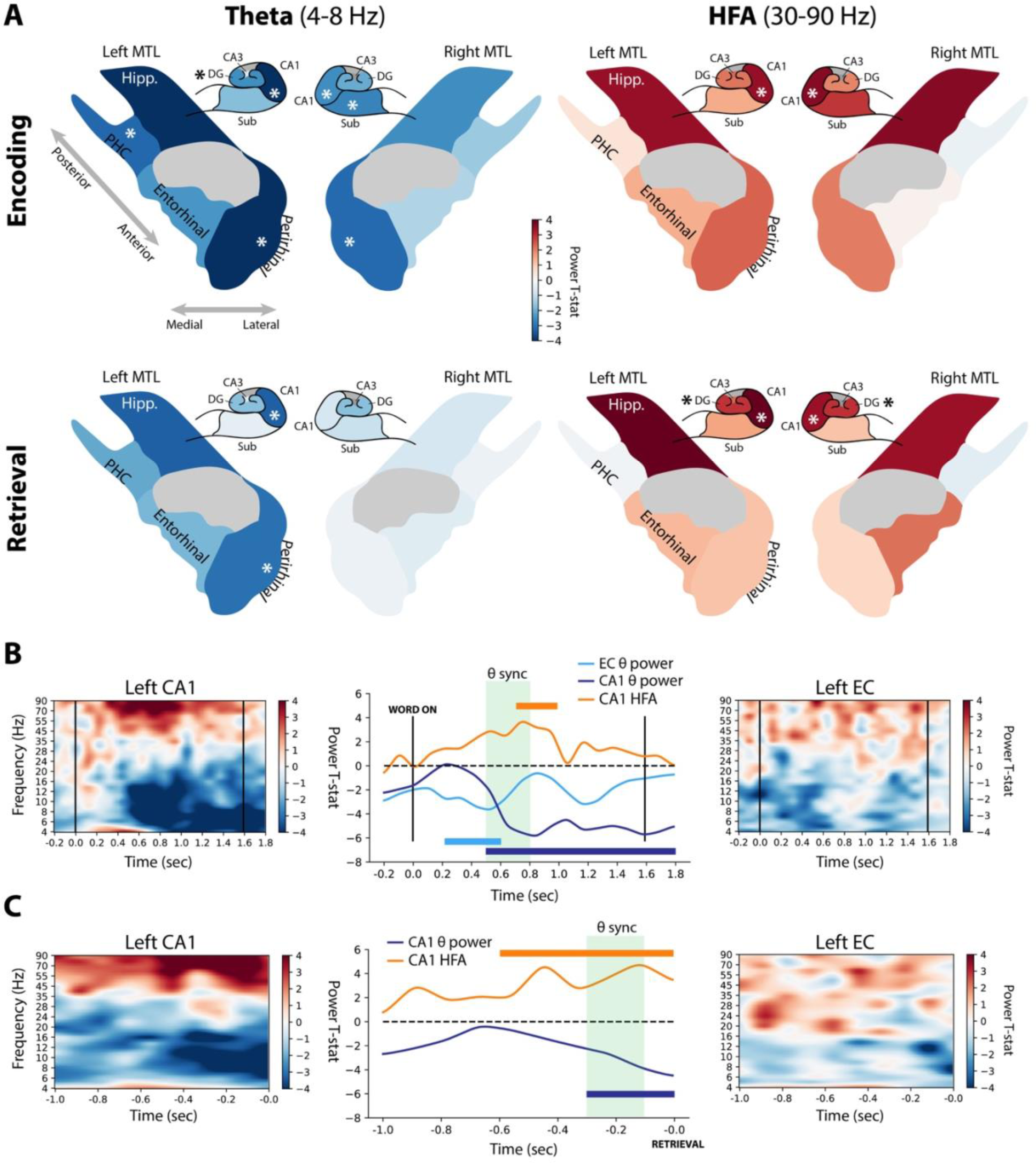
Dynamics of spectral power associated with memory encoding and retrieval. **A.** For each MTL subregion and hippocampal subfield, the spectral power during successful vs. unsuccessful encoding or retrieval epochs was computed in the theta (4-8 Hz) and high-frequency activity (30-90 Hz) bands. For encoding periods, powers were averaged in the 400-1100 ms interval, and between −500-0 ms for retrieval periods, which were the times featuring the most prominent network-wide power change (see Methods for details). The t-statistic indicating the relative power during successful versus unsuccessful encoding or retrieval is mapped to a color, with reds indicating increased power and blues indicating decreased power. These colors are displayed on schematics of MTL and hippocampal anatomy for encoding and retrieval conditions (rows), and theta or HFA bands (columns). Asterisks indicate significant (*P* < 0.05) memory-related power modulation, FDR corrected across tested regions. “Hipp” was not tested collectively but is colored according to CA1. **B.** Left CA1 and left EC showed changes in spectral power that were temporally associated with enhanced connectivity between the regions (see Figure 5B). Significant (*P* < 0.05) increases in CA1 HFA occurred from 700-1000 ms after word onset, while CA1 theta power decreased from 500 ms to the end of the word encoding interval. Left EC theta power decreased from 200-600 ms. The period of significantly enhanced theta PLV is marked in green. **C.** Organized as (B), but for the EC-CA1 interactions in the retrieval contrast. No significant modulations of left EC power were observed in the pre-recall interval.

Between left EC and CA1 – both exhibiting memory-related increases in theta connectivity – we further asked whether there was a relationship between modulations of spectral power and connectivity. During successful encoding, left CA1 showed a significant (*P* < 0.05) increase in HFA from 700-900 ms after word onset, coincident with the 500-800 ms theta connectivity to EC shown in Figure 5B (Figure 6B, top row). Additionally, CA1 exhibited a sustained and significant decrease in theta power beginning at 500 ms, while EC showed a transient decrease from 200 to 600 ms (no significantly increased HFA was observed in EC). In the retrieval contrast, HFA increased and theta power decreased in CA1 prior to onset of a successfully retrieved word (HFA, −600-0 ms prior to onset; theta, −300-0 ms). Both of these intervals overlapped with the period of enhanced CA1-EC theta synchrony from −300 to −100 ms (Figure 6C). We observed no significant modulations of power in either band in EC during retrieval, but note subthreshold increases in HFA and decreases in theta power in the pre-retrieval interval (Figure 6C, right panel). Time-frequency analyses for all MTL regions are reported in Supplemental Figures 2-5. Collectively, these results recapitulate a theme noted in an earlier study of whole-brain connectivity (3): Increases in low-frequency connectivity are associated with increases in high-frequency power and decreases in low-frequency power.

### Memory effects by frequency band

As several frequency bands have been implicated in intra-MTL synchronization (10, 23, 34, 43, 44) – notably theta (4-8 Hz) and low gamma (30-60 Hz) – we finally asked whether memory-related connectivity in the MTL was also present at higher frequency bands. For the theta, alpha (9-13 Hz), beta (16-28 Hz), and low gamma bands, network-wide synchronization was only positive for both encoding/retrieval using PLV in the theta band, though not significantly greater than synchronicity expected by chance (permuted *P* = 0.22, 0.53; Figure 7A; see Methods for details). Specific connection weights for each contrast and connectivity metric are depicted in Figure 7B. Network-wide wPLI was not positive in any band for either contrast. Significant desynchronization was observed for PLV in the beta band and wPLI in the gamma band (permuted *P* < 0.05). In general, synchronization tended to decrease with increasing frequency – considering both PLV and wPLI, and under the bipolar or common average reference, the greatest desynchronization was observed in either the beta or low gamma bands. We note that the bipolar PLV was net positive – though not significant – in the low gamma band, though the effect reversed when considering the bipolar wPLI.

**Figure 7.**
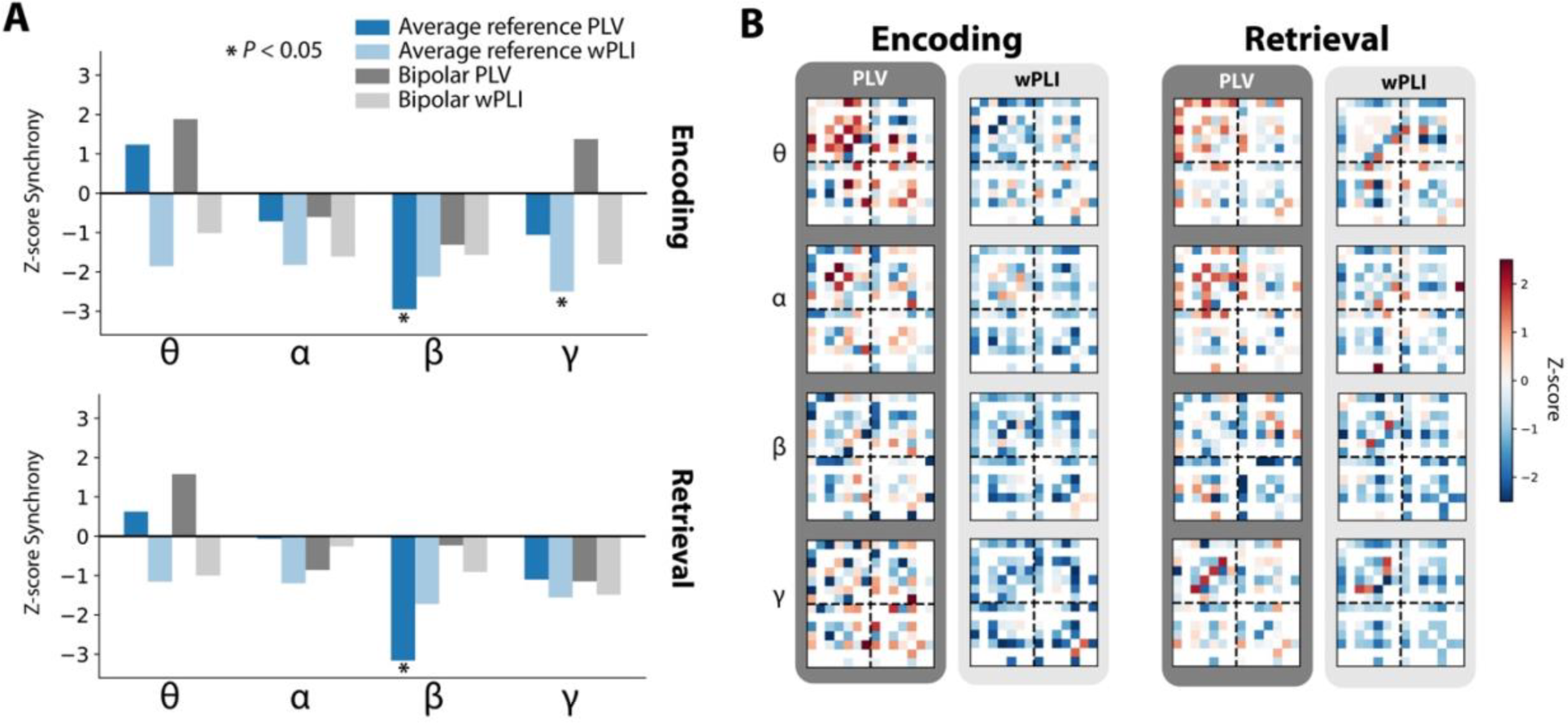
Network-wide synchrony by frequency band. **A.** Network-wide synchronization is computed by averaging all inter-regional connection weights for each frequency, connectivity metric, and referencing scheme. Positive network-wide synchrony was noted for both retrieval and encoding contrasts in the theta band. Significant PLV desynchronization (\**P* < 0.05) was noted in the beta band for encoding and retrieval, and in the gamma band for encoding. B. Adjacency matrices for both contrasts (left vs. right) using PLV and wPLI, organized as shown in Fig. 2A.

Taken together, MTL-wide networks exhibited net increases in connectivity for both encoding and retrieval only in the theta band. These increases were not significantly greater than chance, likely reflecting an underlying mix of synchronous and asynchronous connection weights (Figure 7B). This trend is consistent across referencing scheme and choice of connectivity metric, though strong increases in connectivity using wPLI are not apparent at any band, perhaps due to the reduced sensitivity of this metric if phase differences are near zero. We note that, as shown in earlier analyses, this finding does not preclude the possibility that significantly positive wPLI interactions correlate with successful memory, just that network-wide effects averaged over time could obscure such effects.

## Discussion

We set out to understand neural interactions between substructures of the MTL during episodic encoding and retrieval. As 126 subjects performed a verbal free-recall task, we recorded intracranial EEG from the MTL and compared inter-regional connectivity between periods of successful and unsuccessful memory operations. Using these methods, we discovered that low-frequency phase locking correlates with successful memory encoding and retrieval, with left entorhinal cortex acting as a key hub for theta connectivity during encoding, and a reorganized left-MTL network supporting retrieval. We additionally used the weighted phase-lag index to account for volume conduction, and noted periods of theta synchronization between key regions but generally blunted effects across all frequency bands. Concurrent with these dynamics was a general decrease in theta power and increase in high-frequency activity in both retrieval and encoding, though the degree of power modulation was not predictive of network hubs.

Low-frequency synchronization in the MTL has been conjectured to underlie diverse cognitive operations, including spatial navigation and working memory (10, 17). Under these hypotheses, theta oscillations result in long-term potentiation of synapses in the hippocampus by linking the time of stimulus onset with a cell’s state of maximum depolarization. Relatedly, it is well-established that activity in the gamma-band can be modulated by the phase of an ongoing theta oscillation, representing another mechanism that supports inter-regional plasticity (32, 54, 55). Our data align with this hypothesis – increases in theta connectivity occurred alongside enhanced HFA in connected areas. We show this theta/gamma dynamic in the context of successful encoding of individual words, suggesting that the cognitive operations which support working memory, navigational memory, and episodic memory are not so different; indeed, episodic encoding of list items is known to involve the contextual binding of one item to another (56), not dissimilar to holding a list in working memory or building a map of a spatial environment (20, 57).

Our identification of encoding and retrieval-associated networks enriches computational models of memory in the MTL. An influential theory of MTL function postulates that theta oscillations within the hippocampal-entorhinal system constitute a common substrate of navigation and episodic memory, by synchronizing EC representations of physical or mental space with hippocampal mechanisms that serve to neurally associate these representations with context (and later serve to retrieve information) (18, 19, 30). In support of this theory, we found theta connectivity between the EC and CA1/DG in the hippocampus (in addition to a less robust increase in connectivity between CA1 and DG directly). Enhanced theta connectivity between EC and PRC has not been reported before in humans but supports the notion that EC’s representations are built on sensory input from the neocortex, routed through extrahippocampal MTL regions.

Our findings also indicate a role for theta synchronization during memory retrieval. Indeed, as suggested by anatomical evidence (47) and models of hippocampal function (58), successful retrieval was associated with enhanced connectivity between CA1-EC and CA1-subiculum. Both of these functional connections may support the reinstatement of neocortical activity associated with contextually-retrieved information, driven by pattern completion in CA3.

In this study, we rigorously assessed the utility of two common metrics of phase-based connectivity: the phase-locking value and the weighted phase-lag index. In general, neither metric showed strong evidence for high-frequency synchronization (beta or above), and only the PLV showed truly robust evidence for synchronization in the theta band. Indeed, a total reliance on the wPLI – even in this large dataset – would suggest a general absence of any connectivity-related phenomena during episodic memory processing in the MTL. For several key connections, we demonstrated why there are interpretive difficulties in using the wPLI: it is sensitive to changes in the variance of clustered phase lags *and* the mean direction of clustered phase lags. As such, differences in wPLI between conditions could reflect either (or both) of these underlying dynamics. In regards to the mean direction of phase lags, it is especially difficult to determine whether a change in wPLI is meaningful, because wPLI will statistically discount phase lags as they rotate towards the zero axis, even if they are still well within believable ranges (59). Furthermore, it is not clear that exactly-zero phase lags are biologically implausible and should be ignored outright (6), especially at low frequencies. We observed that, in this dataset, significant increases in PLV were often accompanied by subthreshold increases in wPLI – suggesting that this conservative statistic was, averaged across subjects, reducing physiologic signal to the point that it was no longer significant. At the risk of doubling the number of statistical comparisons, we advise that researchers consider PLV and wPLI simultaneously, and further examine the underlying distribution of phase differences to gain insight into the biological processes that might drive their effects.

Our use of a verbal free-recall task – though a powerful paradigm for studying episodic memory – necessitated the construction of a retrieval contrast that merits further discussion. In this manuscript, we compared neural activity in 1-second intervals leading up to vocalization of a word against 1-second intervals at matched periods of time with no recalls, in free-recall periods from other lists. In this way, we aimed to contrast activity related to successful retrieval against activity during which subjects were liable to try, but fail, to recall a word. This paradigm has been employed in several prior studies examining the neural correlates of free recall (3, 29, 41, 40). However, free-recall tasks inherently confound neural process responsible for episodic retrieval with processes responsible for vocalization and motor preparatory behavior. To account for this, our analyses exclusively consider the MTL – not canonically associated with speech preparation – and only examine activity in the time period preceding onset of vocalization. It is still possible that speech-related activity contaminates the retrieval contrast reported here -- other possible contrasts could leverage nonword vocalizations or intrusion events, though these are typically too rare to serve as a statistically valid basis for connectivity computations. Replicating the finding of recalled-associated theta synchrony in a cued-recall paradigm would therefore be a valuable complement to this work.

In this study, we failed to find strong evidence for memory-related gamma synchronization within the MTL – rather, we noted substantial time-averaged decreases in synchronization in the 30-60 Hz range, especially when using the wPLI. However, earlier reports of hippocampal-rhinal connectivity have reported effects in the gamma band (21, 23). To constrain our hypothesis space, we did not statistically assess possible gamma-band synchronization in detail, though our time-frequency analyses of key hippocampal-rhinal connections such as EC-CA1 and EC-DG do not indicate robust increases in gamma synchronization associated with successful encoding (see Figures 4B and 5B). These data recapitulate an earlier study that utilized a superset of the data here, wherein gamma connectivity was found to decrease during good memory encoding, across a diverse array of cortical regions and hippocampus (3).

In summary, we found that theta band connectivity characterizes intra-MTL interactions that are related to memory encoding and retrieval processes, but distinct networks correlate with successful encoding and retrieval. During encoding, we found EC to be a hub of theta connectivity, with connections to CA1 and DG. During retrieval, we observed a reorganized theta network, with no clear hubs but enhanced connectivity between EC-CA1 and CA1-subiculum. However, both retrieval and encoding were broadly characterized by enhanced HFA and decreased theta power. These findings point to low-frequency interactions as the key to unlocking the way in which medial temporal structures give rise to episodic memories.

## Methods

### Participants

For connectivity analyses, 126 patients with medication-resistant epilepsy underwent a surgical procedure to implant subdural platinum recording contacts on the cortical surface and within brain parenchyma. Contacts were placed so as to best localize epileptic regions. Data reported were collected at 8 hospitals over 3 years (2015-2017): Thomas Jefferson University Hospital (Philadelphia, PA), University of Texas Southwestern Medical Center (Dallas, TX), Emory University Hospital (Atlanta, GA), Dartmouth-Hitchcock Medical Center (Lebanon, NH), Hospital of the University of Pennsylvania (Philadelphia, PA), Mayo Clinic (Rochester, MN), National Institutes of Health (Bethesda, MD), and Columbia University Hospital (New York, NY). Prior to data collection, our research protocol was approved by the Institutional Review Board at participating hospitals, and informed consent was obtained from each participant.

### Free-recall task

Each subject participated in a delayed free-recall task in which they studied a list of words with the intention to commit the items to memory. The task was performed at bedside on a laptop. Analog pulses were sent to available recording channels to enable alignment of experimental events with the recorded iEEG signal.

The recall task consisted of three distinct phases: encoding, delay, and retrieval. During encoding, lists of 12 words were visually presented. Words were selected at random, without replacement, from a pool of high frequency English nouns (http://memory.psych.upenn.edu/WordPools). Word presentation lasted for a duration of 1600 ms, followed by a blank inter-sitmulus interval of 800 to 1200 ms. Before each list, subjects were given a 10-second countdown period during which they passively watch the screen as centrally-placed numbers count down from 10. Presentation of word lists was followed by a 20 second post-encoding delay, during which time subjects performed an arithmetic task during the delay in order to disrupt memory for end-of-list items. Math problems of the form A+B+C=?? were presented to the participant, with values of A, B, and C set to random single digit integers. After the delay, a row of asterisks, accompanied by a 60 Hz auditory tone, was presented for a duration of 300 ms to signal the start of the recall period. Subjects were instructed to recall as many words as possible from the most recent list, in any order, during the 30 second recall period. Vocal responses were digitally recorded and parsed offline using Penn TotalRecall (http://memory.psych.upenn.edu/TotalRecall). Subjects performed up to 25 recall lists in a single session (300 individual words).

### Electrocorticographic recordings

iEEG signal was recorded using depth electrodes (contacts spaced 5-10 mm apart) using recording systems at each clinical site. iEEG systems included DeltaMed XlTek (Natus), Grass Telefactor, and Nihon-Kohden EEG systems. Signals were sampled at 500, 1000, or 1600 Hz, depending on hardware restrictions and considerations of clinical application. Signals recorded at individual electrodes were first referenced to a common contact placed intracranially, on the scalp, or mastoid process. To eliminate potentially confounding large-scale artifacts and noise on the reference channel, we next re-referenced the data using the common average of all depth electrodes in the MTL that were used for later analysis. For some analyses (Figure 2 and Figure 5) raw signals recorded at individual recording contacts were converted to a bipolar montage by computing the difference in signal between adjacent electrode pairs on each depth electrode. Signals were notch filtered at 60 Hz with a fourth-order 2 Hz stop-band butterworth notch filter in order to remove the effects of line noise on the iEEG signal, and downsampled to 256 Hz.

As determined by a clinician, any contacts placed in epileptogenic tissue or exhibiting frequent inter-ictal spiking were excluded from all subsequent analyses. Any subject with fewer than 3 remaining recording contacts in the MTL were not included in the analysis. Any subject with fewer than 15 trials of successful encoding or successful retrieval (see “Retrieval analyses”) were excluded from analysis (encoding, 3 subjects excluded; retrieval, 21 subjects excluded).

#### Limitations of the bipolar reference

In this manuscript, we considered use of the common average and bipolar reference, to account for the possibility that the filtering properties of a given reference scheme could affect connectivity measures. However, the use of the bipolar reference for studies of intra-MTL connectivity is limited by the geometry of linear depth electrodes relative to MTL structures; it is often the case that a bipolar midpoint “virtual” electrode will fall in a subregion/subfield where neither physical contact was placed, raising interpretive difficulties. Additionally, connectivities between bipolar electrodes that share a common monopolar contact are contaminated by shared signal between the two – ideally, such pairs should be excluded from analysis. However, doing so drastically reduces the number of possible region-to-region pairs within the MTL. In the bipolar analyses considered here, all possible pairs were retained even if they shared a common contact, but the bulk of our analyses therefore focus on the average reference. (The use of behavioral contrasts and wPLI may mitigate the effect of shared signal between bipolar virtual contacts.)

### Anatomical localization

To precisely localize MTL depth electrodes, hippocampal subfields and MTL cortices were automatically labeled in a pre-implant, T2-weighted MRI using the automatic segmentation of hippocampal subfields (ASHS) multi-atlas segmentation method (60). Post-implant CT images were coregistered with presurgical T1 and T2 weighted structural scans with Advanced Normalization Tools (61). MTL depth electrodes that were visible on CT scans were then localized within MTL subregions by neuroradiologists with expertise in MTL anatomy. MTL diagrams were adapted with permission from Moore, et al. (62).

### Data analyses and spectral methods

To obtain phase-locking values (PLV) and weighted phase lag index (wPLI) between electrode pairs, we used the MNE Python software package (63), a collection of tools and processing pipelines for analyzing EEG data. PLV reflects the consistency of phase differences between two electrodes across trials (42). wPLI operates similarly to PLV, but weights phase differences according to their rotation away from the zero axis, to account for volume conduction (38). Stated differently, the wPLI weights cross-spectra by the magnitude of the imaginary component of the cross spectrum. Therefore, maximum wPLI is achieved if phase differences are tightly clustered around 90 (or 270) degrees. Both metrics range from 0 (no synchronization) to 1 (maximal synchronization).

To obtain phase information, we convolved signals from each MTL recording contact with compelx-valued Morlet wavelets (6 cycles). We used 24 wavelets from 3-60 Hz as follows: theta (4-8 Hz, spaced 1 Hz), alpha (9-13 Hz, spaced 1 Hz), beta (16-28 Hz, spaced 2 Hz), low gamma (30-60 Hz, spaced 5 Hz). For encoding analyses, each wavelet was convolved with 4000 ms of data surrounding each word presentation (referred to as a “trial”), from 200 ms prior to word onset to 1800 ms afterwards, buffered with 1000 ms on either end (clipped after convolution). Retrieval analyses considered 1000 ms of data prior to each retrieval event, also buffered with 1000 ms on either end.

For each subject, for all possible pairwise combinations of MTL electrodes, we compared the distributions of phase differences in all remembered trials against all not-remembered trials, asking whether there is a significantly higher PLV/wPLI in one or the other. In the encoding contrast, values were compared between all epochs where words were later remembered versus forgotten. In the retrieval contrast, values were compared between epochs leading up to onset of a verbal recall versus matched periods of time when no recall occurred (“deliberation” events, see “Retrieval analysis”). To do this, we found the difference of PLV/wPLI across conditions, e.g.:

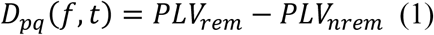

Where *pq* is an electrode pair, *f* is a frequency of interest, and *t* is a window in time. Higher positive differences (*D*) indicate greater connectivity for remembered trials, whereas lower negative differences reflect greater connectivity for not-remembered trials. *D* was computed for each frequency spanning a range from 3 to 60 Hz, averaged into 100 ms non-overlapping windows spanning each trial (i.e. word encoding or pre-retrieval event). 20 windows covered encoding events, from 200 ms prior to word onset to 200 ms after offset. 10 windows covered retrieval/deliberation events, starting 1 second prior to word onset (or 1-second of time during matched deliberation period).

PLV and wPLI values are biased by the number of vectors in a sample. Since our subjects generally forget more words than they remember, we adopt a nonparametric permutation test of significance. For each subject, and each electrode pair, the synchrony computation described above was repeated 250 times with the trial labels shuffled, generating a distribution of *D* statistics that could be expected by chance for every electrode pair, at each frequency and time window. Since only the trial labels are shuffled, the relative size of the surrogate remembered and not-remembered samples also reflect the same sample size bias. Consequently, the true *D* (*D*_true_) can be compared to the distribution of null Ds to derive a z-score or p-value. Higher z-scores indicate greater synchronization between a pair of electrodes for items that are successfully recalled.

To construct a network of synchrony effects between all MTL subregions, we pooled synchrony effects across electrode pairs that span a pair of subregions, and then pooled these subregion-level synchronizations across subjects with that pair of subregions sampled. To do this, we first averaged the *D*_true_ values across all electrode pairs that spanned a given pair of subregions within a subject. Next, we averaged the corresponding null distributions of these electrode pairs, resulting in a single *D*_true_ and a single null distribution for each subregion-pair in a subject. We then averaged the *D*_true_ values and null distributions across all subjects with electrodes in a given ROI pair. By comparing the averaged *D*_true_ to the averaged null distribution, we computed a z-score (and corresponding p-value) at each frequency and temporal epoch that indicates significant synchrony or asynchrony, depending on which tail of the null distribution the true statistic falls.

#### Statistical considerations

Our procedure for averaging the true and null statistics across subjects enables us to construct whole-MTL networks across datasets in which no single subject has electrodes in every region of interest. We compute statistics on these networks that leverage their completeness, including overall connection strength (Figure 2B) and node strengths (Figure 3). Such statistics cannot be assessed at the level of individual subjects who may only have electrode pairs that span a small subset of MTL regions. However, the connection strengths for individual region-pairs can be statistically evaluated across subjects using a 1-sample T-test, so long as a sufficient number of subjects have been sampled for that pair. To demonstrate the correspondence between these two approaches, we correlated the connection weight of population-level z-scores (derived from the permutation procedure above) to t-statistics computed derived from a 1-sample T-test on z-scores from individual subjects. Across all possible region-pairs, connection weights are highly correlated between the two methods (Pearson’r *r* = 0.88, Supplemental Figure 6).

### Network analyses

Using the population-level statistics described above, a 14-by-14 adjacency matrix was constructed for each of the temporal epochs in encoding/retrieval conditions, for each frequency. This matrix represented every possible interaction between all MTL subregions. The z-score of the true *D* relative to the null distribution was used as the connection weight of each edge in the adjacency matrix. Negative weights indicate ROI pairs that, on average, desynchronized when a word was recalled successfully, and positive weights indicate ROI pairs that synchronized when a word was recalled successfully. We zeroed-out any ROI pairs in the matrix represented by less than 5 subjects’ worth of data, to limit the likelihood that our population-level matrix is driven by strong effects in a single or very small number of individuals (see Supplemental Figure 1 for subject counts at each pair).

Since it is possible that collections of weaker connection weights may still account for significant structure in our network, we did not apply a *z*-score threshold before further analyses. To assess for the significance of phenomena at the network level, we instead used 250 null networks that can be constructed on the basis of *D*s derived from the shuffled trial labels to generate a distribution of chance network-level statistics. True statistics were compared to these null distributions to obtain a *P*-value or *z*-score (e.g. network-wide summed connections weights were computed for true and null networks and reported in Fig. 2B).

Adjacency matrices reflect the average connectivity strength during the item presentation interval (0–1600 ms) or retrieval period (−1000-0 ms) for each frequency band. To create them, we averaged true connection strengths within frequency bands, then averaged across all the 100 ms time windows in the encoding/retrieval intervals, and compared the result to the time/frequency average from each of the 250 null networks, resulting in a new Z-score for the time/frequency-averaged network (e.g. Figure 2A).

In analyses of connectivity timecourses (Figures 4-6), intervals are marked as significant so long as the p-value of PLV/wPLI connectivity exceeds a threshold of *P* < 0.05 (relative to the null distribution for that epoch) for at least 2 consecutive 100 ms epochs.

### Hub analysis

To determine which MTL regions act as significant “hubs,” or regions that have enhanced connectivity to many other nodes in the network, we use the node strength statistic from graph theory (Equation 2) (64):

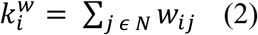

Where *k* is the node strength of node *i*, and *w_ij_* refers to the edge weight between nodes *i* and *j*. *N* is the set of all nodes in the network. In this paper, we only use ipsilateral MTL regions to compute the node strength of each region, so as to (1) better reflect the engagement of a region with its immediate neighbors and (2) acknowledge the sparser sampling of interhemispheric connections. The z-scored connectivity between MTL regions is used as the edge weight. To assess the significance of a hub, we used edge weights derived from each of the 250 null networks, generated by shuffling the original trial labels (see “Network analyses”). For each region, the true node strength is compared to the distribution of null node strengths to derive a z-score or p-value. In Figure 3, p-values were Benjamini-Hochberg corrected for multiple comparisons and thresholded at *P* < 0.05.

### Analysis of spectral power

To determine the change in spectral power associated with successful memory encoding or retrieval, we convolve each electrode’s signal with complex-valued Morlet wavelets (6 cycles) to obtain power information. For high-frequency activity (HFA) we used 13 wavelets spaced 5 Hz (30-90 Hz). Frequencies, time windows, buffers, and spectral methods are otherwise identical to those used in the earlier phase-based analysis (see “Data analyses and spectral methods”).

For each electrode in each subject, we log transformed and z-scored power within each session of the free-recall task, which comprises approximately 300 trials. Power values were next averaged into non-overlapping 100 ms time bins spanning the trial. To assess the statistical relationship between power and later recollection of a word (the power SME), power values for each electrode, trial, time, and frequency were separated into two distributions according to whether the word was later or not remembered, a Welch’s t-test was performed to compare the means of the two distributions. The resulting t-statistics were averaged across electrodes that fell in a common MTL region (either hippocampal subfields or MTL cortices), generating an average t-statistic per subject. Finally, for all MTL regions with more than 5 subjects’ worth of data (all regions except right CA3 met this criteria), we performed a 1-sample t-test on the distribution of t-statistics against zero. The result is a t-statistic that reflects the successful encoding-related change in power across subjects. We report these t-statistics in time-frequency plots in Figure 7B, along with time-averaged t-statistics in Figure 7A (encoding, 400-1100 ms; retrieval, −500-0 ms).

### Retrieval analysis

To find out whether functional connectivity networks uncovered in the memory encoding contrast generalized to different cognitive operations, we further analyzed connectivity in a retrieval contrast. This was done in a manner similar to Burke, et al. 2014 as follows:

For each subject, we identified any 1000 ms period preceding vocal onset of a successfully recalled word, so long as that word was not preceded or followed by any other vocalization for at least 2 seconds. For each retrieval event, we then searched for a 1000 ms interval from a different list during which no successful retrieval (or vocalization) took place, occurring at the same time as the original recall relative to the beginning of the recall period (30-second recall periods followed each of 25 lists per session). These 1000 ms intervals are called “deliberation” intervals, reflecting a time during which a subject was liable to be attempting recall. If no match could be found for the exact time of a given recall, we searched for, still from a different list, a matched deliberation interval within 2 seconds surrounding the onset time of the retrieval event. If no match was available within 2 seconds, the original recall event was discarded from analysis. In this way, each successful retrieval is matched with exactly one deliberation interval, of equal length, from a different recall list.

Analyses of the retrieval contrast were otherwise treated identically to analyses of the encoding contrast, described in “Data analyses and spectral methods.”

## Acknowledgements

We thank Blackrock Microsystems for providing neural recording equipment. This work was supported by the DARPA Restoring Active Memory (RAM) program (Cooperative Agreement N66001-14-2-4032), as well as National Institutes of Health grant MH55687 and T32NS091006. We are indebted to all patients who have selflessly volunteered their time to participate in our study. The views, opinions, and/or findings contained in this material are those of the authors and should not be interpreted as representing the official views or policies of the Department of Defense or the U.S. Government. We also thank Dr. James Kragel for providing valuable feedback on this work.

## Author Contributions

E.S., M.J.K., and D.S.R. designed the study; E.S. analyzed data, and E.S. wrote the paper. J.S., R. Gorniak, S. Das. performed anatomical localization of depth electrodes. M.S., G.W., B.L., C.I., B.J. recruited subjects, collected data, and performed clinical duties associated with data collection including neurosurgical procedures or patient monitoring.

## Data Availability

Raw electrophysiogical data used in this study is freely available at http://memory.psych.upenn.edu/Electrophysiological_Data

## References

1. Eichenbaum H (2000) A cortical-hippocampal system for declarative memory. Nat Rev Neurosci 1(1):41–50.

2. Paller KA, Wagner AD (2002) Observing the transformation of experience into memory. Trends Cogn Sci 6(2):93–102.

3. Solomon EA, et al. (2017) Widespread theta synchrony and high-frequency desynchronization underlies enhanced cognition. Nat Commun 8(1): 1704.

4. Ranganath C, Heller A, Cohen MX, Brozinsky CJ, Rissman J (2005) Functional connectivity with the hippocampus during successful memory formation. Hippocampus 15(8):997–1005.

5. Vincent JL, et al. (2006) Coherent Spontaneous Activity Identifies a Hippocampal-Parietal Memory Network. J Neurophysiol 96(6):3517–3531.

6. Fell J, Axmacher N (2011) The role of phase synchronization in memory processes. Nat Rev Neurosci 12(2):105–118.

7. Eichenbaum H, Otto T, Cohen NJ (1992) The hippocampus--what does it do? Behav Neural Biol 57(1):2–36.

8. Eichenbaum H, Yonelinas AP, Ranganath C (2007) The medial temporal lobe and recognition memory. Annu Rev Neurosci 30:123–52.

9. Wagner AD, et al. (1998) Building memories: remembering and forgetting of verbal experiences as predicted by brain activity. Science 281(5380):1188–91.

10. Buzsáki G (2002) Theta Oscillations in the Hippocampus. Neuron 33(3):325–340.

11. Moser EI, Kropff E, Moser M-B (2008) Place Cells, Grid Cells, and the Brain’s Spatial Representation System. Annu Rev Neurosci 31(1):69–89.

12. MacDonald CJ, Lepage KQ, Eden UT, Eichenbaum H (2011) Hippocampal “time cells” bridge the gap in memory for discontiguous events. Neuron 71(4):737–749.

13. Yamamoto J, Suh J, Takeuchi D, Tonegawa S (2014) Successful Execution of Working Memory Linked to Synchronized High-Frequency Gamma Oscillations. Cell 157(4):845–857.

14. Montgomery SM, Buzsáki G (2007) Gamma oscillations dynamically couple hippocampal CA3 and CA1 regions during memory task performance. Proc Natl Acad Sci U S A 104(36): 14495–500.

15. Schomburg EW, et al. (2014) Theta Phase Segregation of Input-Specific Gamma Patterns in Entorhinal-Hippocampal Networks. Neuron 84(2):470–485.

16. Seidenbecher T, Laxmi TR, Stork O, Pape H-C (2003) Amygdalar and hippocampal theta rhythm synchronization during fear memory retrieval. Science 301(5634):846–50.

17. Colgin LL (2016) Rhythms of the hippocampal network. Nat Rev Neurosci 17(4). 10.1038/nrn.2016.21.

18. Hasselmo ME, Eichenbaum H (2005) Hippocampal mechanisms for the context-dependent retrieval of episodes. Neural Netw 18(9):1172–90.

19. Diana RA, Yonelinas AP, Ranganath C (2007) Imaging recollection and familiarity in the medial temporal lobe: a three-component model. Trends Cogn Sci 11(9):379–386.

20. Burgess N, Maguire EA, O’Keefe J (2002) The Human Hippocampus and Spatial and Episodic Memory. Neuron 35(4):625–641.

21. Fell J, et al. (2001) Human memory formation is accompanied by rhinal-hippocampal coupling and decoupling. Nat Neurosci 4(12):1259–64.

22. Fell J, Ludowig E, Rosburg T, Axmacher N, Elger CE (2008) Phase-locking within human mediotemporal lobe predicts memory formation. Neuroimage 43(2):410–419.

23. Fell J, et al. (2003) Rhinal-hippocampal theta coherence during declarative memory formation: interaction with gamma synchronization? Eur J Neurosci 17(5):1082–1088.

24. Lega BC, Jacobs J, Kahana M (2012) Human hippocampal theta oscillations and the formation of episodic memories. Hippocampus 22(4):748–761.

25. Kahana MJ, Seelig D, Madsen JR (2001) Theta returns. Curr Opin Neurobiol 11(6):739–744.

26. Hasselmo ME (2005) What is the function of hippocampal theta rhythm?--Linking behavioral data to phasic properties of field potential and unit recording data. Hippocampus 15(7):936–49.

27. Lin J-J, et al. (2017) Theta band power increases in the posterior hippocampus predict successful episodic memory encoding in humans. Hippocampus 27(10):1040–1053.

28. Greenberg JA, Burke JF, Haque R, Kahana MJ, Zaghloul KA (2015) Decreases in theta and increases in high frequency activity underlie associative memory encoding. Neuroimage 114:257–63.

29. Burke JF, et al. (2014) Theta and high-frequency activity mark spontaneous recall of episodic memories. J Neurosci 34(34):11355–65.

30. Buzsáki G, Moser EI (2013) Memory, navigation and theta rhythm in the hippocampal-entorhinal system. Nat Neurosci 16(2):130–138.

31. Buzsáki G, Schomburg EW (2015) What does gamma coherence tell us about interregional neural communication? Nat Neurosci 18(4):484–489.

32. Rutishauser U, Ross IB, Mamelak AN, Schuman EM (2010) Human memory strength is predicted by theta-frequency phase-locking of single neurons. Nature 464(7290):903–907.

33. Watrous AJ, Tandon N, Conner CR, Pieters T, Ekstrom AD (2013) Frequency-specific network connectivity increases underlie accurate spatiotemporal memory retrieval. Nat Neurosci 16(3):349–56.

34. Colgin LL (2013) Mechanisms and Functions of Theta Rhythms. Annu Rev Neurosci 36(1):295–312.

35. Sarnthein J, Petsche H, Rappelsberger P, Shaw GL, von Stein A (1998) Synchronization between prefrontal and posterior association cortex during human working memory. Proc Natl Acad Sci U S A 95(12):7092–6.

36. Clouter A, Shapiro KL, Hanslmayr S (2017) Theta Phase Synchronization Is the Glue that Binds Human Associative Memory. Curr Biol 27(20):3143–3148.e6.

37. Backus AR, Schoffelen JM, Szebényi S, Hanslmayr S, Doeller CF (2016) Hippocampal-prefrontal theta oscillations support memory integration. Curr Biol 26(4). 10.1016/j.cub.2015.12.048.

38. Vinck M, Oostenveld R, van Wingerden M, Battaglia F, Pennartz CMA (2011) An improved index of phase-synchronization for electrophysiological data in the presence of volume-conduction, noise and sample-size bias. Neuroimage 55(4):1548–1565.

39. Burke JF, et al. (2014) Human intracranial high-frequency activity maps episodic memory formation in space and time. Neuroimage 85:834–843.

40. Kragel JE, et al. (2017) Similar patterns of neural activity predict memory function during encoding and retrieval. Neuroimage 155:60–71.

41. Long NM, et al. (2017) Contextually Mediated Spontaneous Retrieval Is Specific to the Hippocampus. Curr Biol 27(7):1074–1079.

42. Lachaux JP, Rodriguez E, Martinerie J, Varela FJ (1999) Measuring phase synchrony in brain signals. Hum Brain Mapp 8(4):194–208.

43. Igarashi KM, Lu L, Colgin LL, Moser M-B, Moser EI (2014) Coordination of entorhinal-hippocampal ensemble activity during associative learning. Nature 510(7503):143–147.

44. Fell J, et al. (2001) Human memory formation is accompanied by rhinal-hippocampal coupling and decoupling. 10.1038/nn759.

45. Squire LR, Stark CEL, Clark RE (2004) THE MEDIAL TEMPORAL LOBE. Annu Rev Neurosci 27(1):279–306.

46. Davachi L (2006) Item, context and relational episodic encoding in humans. Curr Opin Neurobiol 16(6):693–700.

47. Kloosterman F, Witter MP, Van Haeften T (2003) Topographical and laminar organization of subicular projections to the parahippocampal region of the rat. J Comp Neurol 455(2):156–171.

48. Liang JC, Preston AR (2015) Medial Temporal Lobe Subregional Function in Human Episodic Memory. The Wiley Handbook on the Cognitive Neuroscience of Memory (John Wiley & Sons, Ltd, Chichester, UK), pp 108–130.

49. Holsheimer J, Feenstra BW (1977) Volume conduction and EEG measurements within the brain: a quantitative approach to the influence of electrical spread on the linear relationship of activity measured at different locations. Electroencephalogr Clin Neurophysiol 43(1):52–8.

50. Diana RA, Yonelinas AP, Ranganath C (2010) Medial temporal lobe activity during source retrieval reflects information type, not memory strength. J Cogn Neurosci 22(8):1808–18.

51. Merkow MB, Burke JF, Kahana MJ (2015) The human hippocampus contributes to both the recollection and familiarity components of recognition memory. Proc Natl Acad Sci U S A 112(46):14378–83.

52. Burke JF, et al. (2015) Human intracranial high-frequency activity during memory processing: neural oscillations or stochastic volatility? This review comes from a themed issue on Brain rhythms and dynamic coordination. Curr Opin Neurobiol 31:104–110.

53. Burke JF, et al. (2013) Synchronous and asynchronous theta and gamma activity during episodic memory formation. J Neurosci 33(1):292–304.

54. Lisman JE, Jensen O (2013) The Theta-Gamma Neural Code. Neuron 77(6):1002–1016.

55. Canolty RT, et al. (2006) High Gamma Power Is Phase-Locked to Theta Oscillations in Human Neocortex. Science (80-) 313(5793):1626–1628.

56. Howard MW, Kahana MJ (2002) A Distributed Representation of Temporal Context. J Math Psychol 46(3):269–299.

57. Howard MW, Fotedar MS, Datey A V., Hasselmo ME (2005) The Temporal Context Model in Spatial Navigation and Relational Learning: Toward a Common Explanation of Medial Temporal Lobe Function Across Domains. Psychol Rev 112(1):75–116.

58. Hasselmo ME, Bodelón C, Wyble BP (2002) A Proposed Function for Hippocampal Theta Rhythm: Separate Phases of Encoding and Retrieval Enhance Reversal of Prior Learning. Neural Comput 14(4):793–817.

59. Cohen MX (2015) Effects of time lag and frequency matching on phase-based connectivity. J Neurosci Methods 250:137–146.

60. Yushkevich PA, et al. (2015) Automated volumetry and regional thickness analysis of hippocampal subfields and medial temporal cortical structures in mild cognitive impairment. Hum Brain Mapp 36(1):258–87.

61. Avants BB, Epstein CL, Grossman M, Gee JC (2008) Symmetric diffeomorphic image registration with cross-correlation: Evaluating automated labeling of elderly and neurodegenerative brain. Med Image Anal 12(1):26–41.

62. Moore M, et al. (2014) A comprehensive protocol for manual segmentation of the medial temporal lobe structures. J Vis Exp (89). 10.3791/50991.

63. Gramfort A, et al. (2014) MNE software for processing MEG and EEG data. Neuroimage 86:446–460.

64. Rubinov M, Sporns O (2010) Complex network measures of brain connectivity: Uses and interpretations. Neuroimage 52(3):1059–1069.

